# *In virio* secondary RNA structure analysis of influenza A virus

**DOI:** 10.64898/2026.02.09.704913

**Authors:** Astha Joshi, Lin-Chen Huang, Adrianus C. M. Boon

## Abstract

The genome of influenza A viruses (IAV) consists of eight distinct gene segments that need to be packaged into new virions. Although this process is thought to be controlled by specific RNA-RNA interactions between segments, few studies have identified the RNA structure inside virions and performed functional validations on these structures. Using high-throughput probing techniques such as SHAPE-MaP and DMS-MaPseq, we mapped the secondary structure of the A/Puerto Rico/8/1934 (PR8) IAV genome. We discovered 173 putative structural motifs in both packaging and central regions of all segments. Fifteen motifs were selected for functional studies by introducing synonymous mutations to destabilize the predicted structure and assessing viral fitness. Synonymous mutations in three predicted RNA structures within the packaging signals of NP and PB1 attenuated virus replication *in vitro*. Interestingly, combining structural changes in multiple motifs within the packaging signals enhanced viral attenuation, supporting the idea of synergistic structural contributions to viral fitness. In contrast, structural synonymous mutations in RNA secondary structures outside packaging signals had no impact on virus replication *in vitro*. Overall, our findings revealed several new RNA secondary structures within the packaging signals of IAV gene segments that play a critical role in the IAV life cycle.

## INTRODUCTION

The influenza A virus (IAV) poses a significant public health risk due to its potential to cause epidemics, pandemics, and zoonotic outbreaks, resulting in substantial societal and economic burdens each year (1, 2). One notable historical example is the devastating Spanish flu pandemic in 1918, which resulted in the deaths of millions worldwide (3). According to the World Health Organization (WHO), annual influenza epidemics lead to three to five million cases of severe illness globally and up to half a million deaths, particularly among high-risk populations such as children, the elderly, and immunocompromised individuals. This impact is particularly severe in developing countries.

Influenza A virus belongs to the *Orthomyxoviridae* family (4). It is an enveloped virus with a negative-strand RNA genome consisting of eight gene segments - PB2, PB1, PA, HA, NP, NA, M, and NS - which encode a minimum of 14 viral proteins. PB2, PB1, and PA together form a heterotrimeric polymerase complex essential for viral replication and the transcription of viral RNA (5, 6). The polymerase complex operates within the nucleus of infected cells, where it interacts with viral RNA (vRNA) and cellular factors to carry out its functions (7). Each vRNA contains a central coding region flanked by 5’ and 3’ non-coding regions (NCRs). The terminal regions contain complementary sequences allowing the formation of a secondary RNA structure that enables polymerase binding and activity (5). The vRNA, the nucleoprotein (NP), and the polymerase complex constitute the complete viral ribonucleoprotein complex (vRNP) (8, 9). For IAV to be fully infectious, its virions must contain all eight gene segments (10). The prevailing IAV model suggests a selective packaging process, where the eight different vRNPs are chosen hierarchically (11). This selection is facilitated by distinct packaging signals found in the terminal regions of each vRNA, allowing discrimination among them (12, 13). Packaging signals are specific cis-acting RNA elements on each segment that ensure selective incorporation into new virions during viral assembly. They were identified through mutagenesis studies, reporter assays, and bioinformatic analysis (12, 14–17) and vary in length based on the gene segment. Each vRNP interacts with at least one other vRNP partner to form a larger complex (18), likely maintained by intra-or inter-RNA-RNA interactions between different RNA segments or between RNA and proteins, which are thought to guide the packaging process. However, the precise mechanism governing the selection, interaction, and arrangement remains poorly understood.

The intricate interplay between RNA structure and function is pivotal in the viral life cycle. RNA structure probing techniques, combined with high-throughput sequencing, enable the detection of structural characteristics of RNA molecules across the entire transcriptome. These methods offer valuable insights into the previously underestimated role of RNA structure within the intricate cellular environment. The significance of RNA structure in regulating and fine-tuning cellular processes has become especially evident with the application of transcriptome-wide structure probing techniques (19). Additionally, mutational profiling (MaP) development has enhanced the feasibility of RNA structure probing methods to address pertinent biological inquiries. MaP increases coverage across transcripts by allowing read-through at probing sites, thus eliminating the need for size-selection of truncated cDNA fragments. A recent study on different viruses using these high-throughput probing techniques shows that viral RNAs can form a stable and conserved structure inside a cell that plays critical roles in replication, transcription, packaging, and host interaction (20–22). Disruption of these structures leads to the attenuation of the viral replication process.

Although many studies have identified putative RNA secondary structures, as well as intra- or inter RNA-RNA interactions, very few have demonstrated what their significance may be (14–17, 23). Three intra-segment structures have been functionally studied in segments 1, 4, and 5. The PSL2 motif RNA structure was identified in the 5’ packaging signal of segment 1 (PB2), stem-loop structure in the 3’ packaging signal of segment 4 (HA), and the pseudoknot structure in segment 5 (NP). Synonymous mutations intended to disrupt the predicted RNA secondary or tertiary structures reduced viral fitness and affected genome packaging (24, 25–30). These essential RNA secondary structures near the 3’ and 5’ ends of gene segments, support the prevailing model of IAV genome packaging.

In this study, we used SHAPE-MaP (selective 2′-hydroxyl acylation analyzed by primer extension and mutational profiling) and DMS-MaPseq (dimethyl sulfate (DMS) mutational profiling with sequencing) (31, 32) to identify RNA secondary structures in the genome of an H1N1 IAV. To assess the significance of these predicted RNA secondary structures, synonymous structural changes were introduced, and viral fitness was assessed.

## MATERIALS AND METHODS

### Cell culture and IAV propagation and purification

Madin–Darby canine kidney (MDCK) cells were maintained in minimal essential medium (MEM) with 10% fetal bovine serum (FBS, Biowest), 1x MEM vitamins (Gibco), 2 mM L-glutamine (Gibco), and 100 U/mL Penicillin and 100 µg/mL Streptomycin (Gibco). Human embryonic kidney cells (293T) were maintained in Opti-MEM (Life Technologies) with 10% FBS, 2 mM L-glutamine, and 100 U/mL Penicillin and 100 µg/mL streptomycin. MDCK and 293T cells were a kind gift from Dr. Richard Webby at St. Jude Children’s Research Hospital. Unless indicated, all cell cultures were incubated at 37°C in 5% CO2.

Stocks of A/Puerto Rico/8/1934 (PR8, H1N1) virus were produced by either infecting MDCK cells or 10-day-old embryonated chicken eggs. The MDCK cells were grown to confluency and infected with IAV in MEM media containing 0.1% BSA at a multiplicity of infection (MOI) of 0.001. After 1 hour, the media containing the virus was removed, and a new media, supplemented with 1 μg/mL of n-Tosyl-l-phenylalanine chloromethyl ketone (TPCK)-treated trypsin (Merck) was added. Two days later, the supernatant was collected, aliquoted, and stored at −80°C. PR8 was also grown in 10-day old embryonated chicken eggs at 37°C. Two days after inoculation, the allantoic fluid was collected, clarified by centrifugation at 2000x*g* for 20 minutes, and stored at −80°C. Virus containing cell culture supernatant or allantoic fluid was treated with RNase A and RNase III at 37°C for an hour to get rid of exogenous RNA. Next, the supernatant or allantoic fluid was clarified by centrifugation at 3,200x*g* for 10 min at 4°C, followed by centrifugation at 16,000x*g* for 15 min at 4°C. Next, the fluid was mixed with 40% PEG 6000 (Sigma, catalog #528877) at 10% final PEG-6000 concentration and left overnight on a shaker at 4°C. The next day, the sample was spun at 1,600x*g* for 60 min at 4°C, the supernatant was carefully removed, and the purified virus pellet was resuspended in a buffer (0.05 M HEPES (pH 8), 0.1 M NaCl, 0.0001 M EDTA for SHAPE-MaP or 0.3 M HEPES (pH 8), 0.1 M NaCl for DMS-MaPseq).

### SHAPE-MaP

For selective hydroxyl acylation analyzed by primer extension (SHAPE) modification, the resuspended virus was divided into 3 reactions (modified sample, control sample, and denatured sample). For the modified sample, 1-methyl-7-nitroisatoicanhydride (1M7) (33) or 2-methylnicotinic acid imidazolide (NAI) (19) was added at a final concentration of 100 mM. For the control sample, the corresponding amount of DMSO was added. Samples were then incubated at 37°C for 5 min (1M7) or 15 min (NAI), followed by quenching of NAI through the addition of dithiothreitol (DTT) at a final concentration of 0.5 M. For the denatured sample, the resuspended virus was set aside without any treatment. TRK lysis buffer was added to the above reactions to lyse the virus. Total RNA was extracted using Zymo RNA Clean and Concentrator-5 Kit (Zymo Research). The RNA from the denatured sample was incubated at 95°C for 1 minute and then treated with 100 mM 1M7 or NAI for 1 minute at 95°C, and the reaction was quenched with DTT as described previously. For the denatured sample, RNA was again purified using Zymo RNA Clean and Concentrator-5 Kit (Zymo Research). Sequencing library preparation was performed according to the amplicon workflow as described previously (19, 34). The extracted or re-extracted RNA was reverse-transcribed by incubating at 25°C for 10 minutes and 42°C for 180 minutes using random primers together with SuperScript II (Invitrogen) in MaP buffer (50 mM Tris-HCl (pH 8.0), 75 mM KCl, 6 mM MnCl_2_, 10 mM DTT, and 0.5 mM deoxynucleoside triphosphate). The reaction was heat-inactivated at 70°C for 15 minutes, and cDNA was purified. NEBNext^®^ Ultra™ II Non-Directional RNA Second Strand Synthesis Module (New England Biolabs) was used to generate double-stranded cDNA by incubating at 16°C for 150 minutes. The Nextera XT DNA Library Preparation Kit (Illumina) was used to prepare the sequencing libraries. Final PCR amplification products were size selected using Agencourt AMPure XP Beads (Beckman Coulter). Libraries were quantified using a Qubit dsDNA HS Assay Kit (ThermoFisher, Cat. No. Q32851) to determine the DNA concentration, and quality was assessed with the Agilent High Sensitivity DNA kit (Agilent Technologies) on a Bioanalyzer 2100 System (Agilent Technologies) to determine average library member size and accurate concentration. The libraries were sequenced (2 × 150 base pairs (bp)) on a MiniSeq System (Illumina). Sequencing reads were aligned with the reference sequences, and SHAPE-MaP reactivity profiles for each position were calculated using Shapemapper-2.15 with the min-depth option set to 1000 (35). All SHAPE-MaP reactivities were normalized to an approximate 0–2 scale by dividing the SHAPE-MaP reactivity values by the mean reactivity of the 10% most highly reactive nucleotides after excluding outliers (defined as nucleotides with reactivity values that are >1.5 the interquartile range). High SHAPE-MaP reactivities above 0.7 indicate more flexible (that is, single-stranded) regions of RNA, and low SHAPE-MaP reactivities below 0.3 indicate more structurally constrained (that is, base-paired) regions of RNA.

### DMS-MaPseq

For di-methyl sulfate (DMS) modification, the resuspended virus was divided into 2 reactions (modified and control sample). For the modified sample, 0.5% v/v DMS (Sigma, catalog #D186309) was added, mixed thoroughly, and incubated immediately at 37° C for 5 min before quenching with 100 µL 30% β-mercaptoethanol in PBS. The control sample was prepared similarly, only without the addition of DMS. TRK lysis buffer was added to the above reactions to lyse the virus. Total RNA was extracted using Zymo RNA Clean and Concentrator-5 Kit (Zymo Research). For reverse transcription, the 11.5 µL RNA was supplemented with 4 μL 5× first strand buffer (ThermoFisher Scientific), 1 μL 10 μM specific reverse primer mix, 1 μL dNTP, 1 μL 0.1 M DTT, 1 μL RNaseOUT (Invitrogen), and 0.5 μL MarathonRT (Kerafast, Catalog #EYU007). The reverse-transcription reaction was incubated at 42° C for 3 hours. 1 μL RNase H was added to each reaction and incubated at 37°C for 20 min to degrade the RNA. cDNA was purified using the QIAquick PCR Purification Kit (Cat. No. 28104). cDNA was converted to dsDNA using Q5 polymerase with segment-specific primers (**Supplementary Table 1**). The library preparation, cleanup, quantification, and sequencing of libraries were performed as described above. DMS-MaPseq reactivity profiles are calculated using the DREEM Webserver, as described previously (34, 36). Also, using ‘Shapemapper-2.15’ with DMS option and with the min-depth option set to 1000.

### RNA structure predictions

SHAPE-MaP and DMS-MaPseq reactivity data obtained through Shapemapper-2.15 were used as constraints to predict the RNA secondary structure using Fold in RNAstructure software suite (v5.8.1) (37) with default settings and md (maximum pairing distance) option set to 150. ShapeKnots (38) in the RNAstructure software suite (v5.8.1) was used to verify the pseudoknot structure present in segment 5. Consensus secondary structure, base pairing probabilities, and Shannon entropies at each position were obtained through Superfold v1.0 (31). Regional entropies were generated by finding the median Shannon entropy over a 50-nucleotide rolling window. Highly structured regions were defined as regions with low median Shannon entropy and low SHAPE reactivity.

### Selection and functional validation of RNA secondary structures in PR8 virus

A total of 15 predicted RNA secondary structures in the PB2, PB1, PA, and NP gene segments were selected for mutational analysis. The predicted RNA secondary structures in the canonical packaging signals of gene segments (n = 6) and the more central open reading frame (n = 9) were included. The selection was based on an analysis of open reading frames to assess potential nucleotide degeneracy while retaining the coding sequence. Synonymous nucleotide changes, predicted to disrupt the vRNA secondary structure but not alter the protein sequence, were introduced into pHW2000 bidirectional plasmids using inverse PCR with primers containing the corresponding alterations. All plasmid constructs were verified by Sanger Sequencing (Genewiz) to confirm the presence of the mutation of interest and the absence of unwanted mutations. Wild-type or mutant PR8 viruses were generated by the transfection of eight plasmids, representing all eight segments of PR8 IAV, into co-cultures of 293T and MDCK cells (1 μg per plasmid) with 18 µL TransIT®-LT1 (Mirus, Cat# MIR 2304) transfection reagent in Opti-MEM. The next day, the transfection mixture was removed and replaced with Opti-MEM containing 0.2% BSA and 1 µg/mL TPCK-trypsin. Seventy-two hours after the addition of TPCK-trypsin, culture supernatant was collected and clarified by centrifugation for 5 min at 350x*g*. The virus titers in the tissue culture supernatant were determined by plaque assays on MDCK cells. Viral stocks were generated by infection of MDCK cells with a fixed MOI of 0.001 and collection of the supernatant 48 hours later. Stocks were aliquoted and stored at −80°C until use. All studies were conducted with passage 1 stocks following sequence verification of the mutation. All viruses were generated at least twice independently.

### Plaque assay

Twenty-four well plates containing confluent monolayers of MDCK cells were inoculated with 250 µL of 10-fold serially diluted virus stock or culture supernatant for 1 h in MEM supplemented with 0.1% BSA and 1 µg/mL TPCK-trypsin. After 1 h, the inoculum was removed and replaced with an overlay of 1% Low Melting Point Agarose (Stellar Scientific) in MEM supplemented with 0.1% BSA and 1 µg/mL TPCK-trypsin. Seventy-two hours post-inoculation, the cells were fixed with 5% formaldehyde for one hour. The agarose plug was then removed, and cells were stained for 1 hour with 1% crystal violet in PBS. Crystal violet was removed, and cells were carefully washed with water. Plaques were counted manually, and virus titer (plaque-forming units (PFU)*/*mL) was calculated.

### Determination of viral replication kinetics

The replication characteristics of wild-type and mutant PR8 viruses were determined by infecting MDCK cells at an MOI of 0.01. Following 1 h absorption (at *t* = 0 h) the inoculum was removed, the cells were washed once with PBS, and fresh 2.0 mL of infection medium was added to the cells. Cell culture supernatants were collected at 24, 32, and 48 h post-inoculation and stored at −80°C for titration by plaque assay.

### Mini-genome luciferase assay with polymerase genes of influenza A virus

The PB2, PB1, PA, and NP genes of PR8 were cloned into pcDNA3.1+ expression vectors. The firefly luciferase gene was cloned into the pLuci vector flanked by the wild-type or mutant 3’ and 5’ untranslated regions (UTRs) from the PR8 PB1 segment. A 100 ng of each plasmid pcDNA3.1-PR8-PB2, PB1, PA, and NP, together with the firefly reporter construct and the Renilla luciferase internal control, was transfected into 293T cells (2.5 × 10 cells/well) in 96-well plates using TransIT®-LT1. After 24 hours, the amount of firefly and Renilla luciferase activity was quantified with the Dual luciferase assay reporter system (Promega, Cat# E1960). Each condition (set of plasmids) was done in triplicate and repeated independently in separate experiments at least three times. The relative light units (RLU) of Firefly are normalized to the RLU for Renilla, and the activity of varying polymerase combinations are then normalized to the positive control. Statistical analysis was done on the average value from a single assay.

### Gene-segment specific qPCR assay

RNA was extracted from wildtype or mutant PR8 viruses (P1 stock) following lysis in 300 μL of TRK lysis buffer, using Zymo RNA Clean and Concentrator-5 Kit (Zymo Research). The RNA was eluted in 20 μL DEPC H_2_O (RNA Clean & Concentrator, Zymo). cDNA was synthesized from 10 μL of RNA with SSIII Reverse transcriptase (Invitrogen) and a vRNA-specific primer (39). Total cDNA was diluted 1:50 and used to quantify each of the 8 genome segments by SYBR Green qPCR (PowerUpTM SYBR® Green Master Mix) with primer pairs previously published (39). Relative abundance of each genome segment was calculated as before, except with normalization to Segment 7.

### Statistical analysis

The data was graphed using GraphPad Prism 10. For comparison of SHAPE-MaP data sets, we used the Python-based Pearson correlation coefficient function (corr()). Two-way ANOVA with multiple comparison corrections (Dunnett’s and Tukey’s test) was used to assess significant differences in virus titer and polymerase activity. We expressed the data as the geometric mean ± standard deviation. *p-values* of ≤ 0.05 were considered significant.

### Data repository

The FASTA files for SHAPE-MaP and DMS-MaPseq can be found under BioProject accession no. PRJNA1403019. All other data supporting the findings of this study are available within the article and its Supplementary Information files or are available from the authors upon request.

## RESULTS

### High-throughput probing analysis enables in virio RNA secondary structure prediction of the influenza A virus genome

Studying the molecular details of vRNA structure and vRNA-vRNA interactions is crucial for understanding the process of viral replication and genome packaging. To predict the vRNA structure of the IAV genome, we performed extensive RNA secondary structure analysis on purified A/Puerto Rico/8/1934 (PR8, H1N1) virus from eggs or cells using SHAPE-MaP and DMS-MaPseq (**Fig 1**). The viral supernatant collected from cells or eggs was treated with RNase A and H to get rid of exogenous RNAs. Both techniques provide information about individual base reactivity at single-nucleotide resolution, which in turn indicates the nucleotide’s relative flexibility that can be correlated with base-pairing likelihood. Nucleotides with a reactivity above 0.6 (DMS-MaPseq) or 0.7 (SHAPE-MaP) are likely unpaired, which means these bases don’t interact with either RNA or protein.

**Figure 1.**
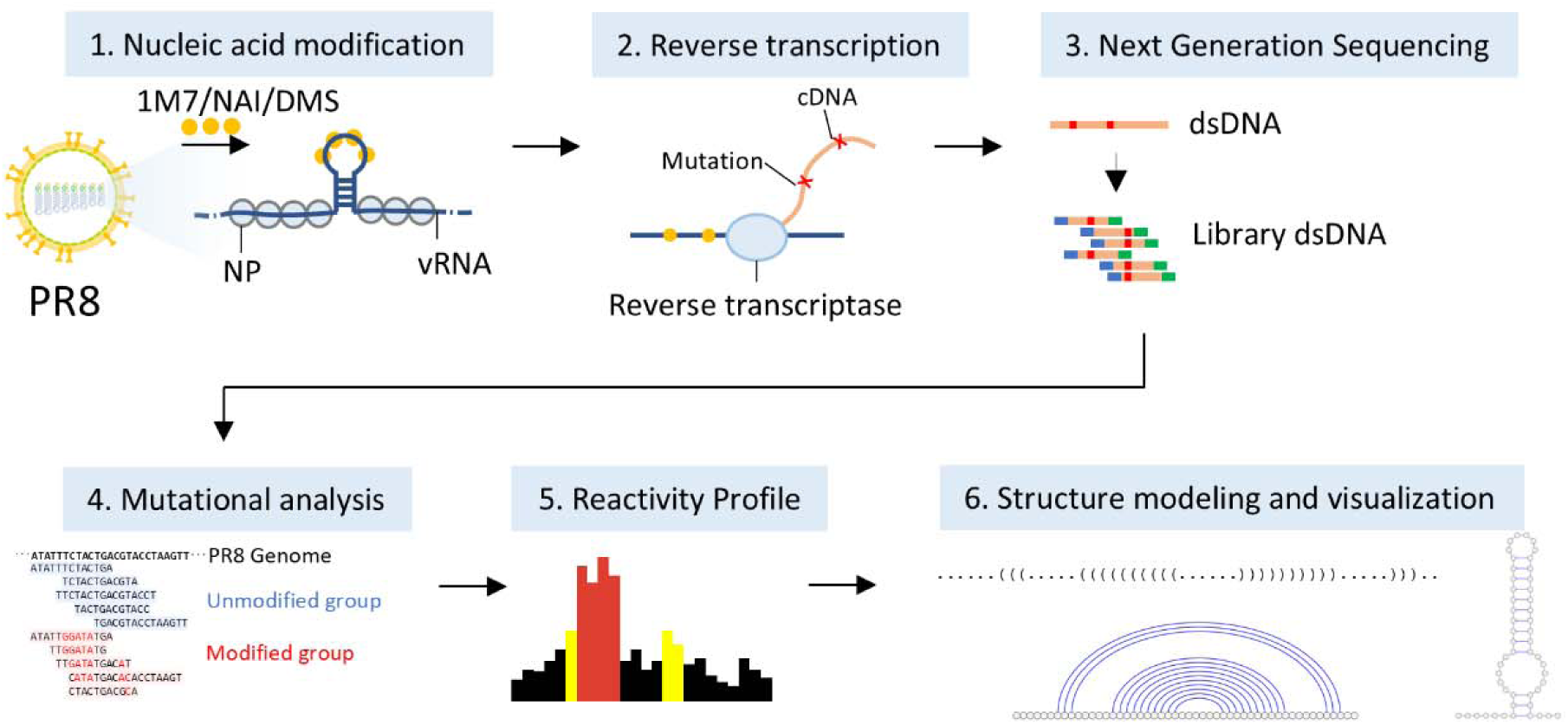
Schematic of the experimental workflow for SHAPE-MaP and DMS-MaPseq. (**1**) Nucleic acid modification in the viral RNA (vRNA) genome inside concentrated influenza A virus particles using SHAPE reagent (1M7 or NAI), or DMS. (**2**) Reverse transcription primer extension reaction on vRNA extracted from control or treated viral samples. (**3**) Next-generation sequencing libraries were prepared using the Nextera DNA library kit and sequenced 2×150bp. (**4**) Sequences were aligned to the reference genome, and mutation rates were calculated. (**5**) Mutation rates were converted to SHAPE reactivity values. (**6**) Secondary structure predictions using SHAPE reactivity as a pseudo energy constraint and different ways of visualizing secondary structure. PR8 (A/Puerto Rico/8/1934), 1M7 (1-methyl-7-nitroisatoic anhydride), NAI (2-methylnicotinic acid imidazolide), DMS (dimethyl sulphate), vRNA (viral RNA), dsDNA (double-stranded DNA).

To establish the pipeline, we first compared different purification methods and RNA-probing reagents. SHAPE-MaP analysis with 1M7 as a probing reagent on PR8 virus that was purified by ultracentrifugation or PEG-precipitation showed a high degree of similarity in nucleotide reactivity with correlations ranging from 0.88-0.93 across different gene segments. Next, we compared two RNA-probing reagents, 1M7 and NAI. Both reagents yielded highly reproducible results; different replicates of 1M7 yielded a high degree of correlation (0.79-0.93), as did different replicates of NAI (0.61-0.81) across all eight gene segments (**Fig S1A**). Overall, the reactivity profile obtained using 1M7 and NAI shows a similar pattern of reactivity (**Fig 2**) and good correlation ranging from 0.58 to 0.82 across different segments (**Fig S1B**).

**Figure 2.**
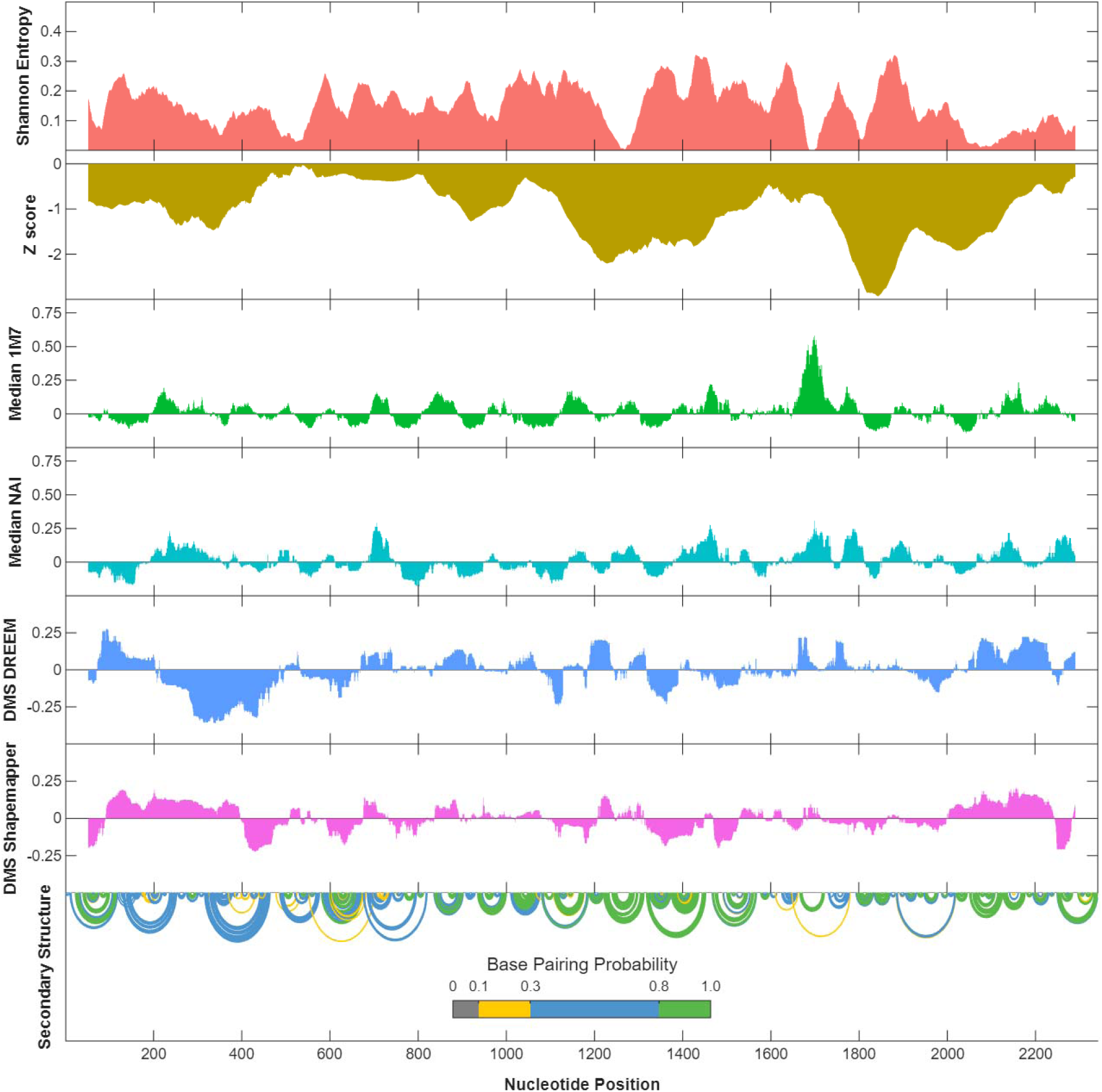
Genome-wide *in virio* secondary structure model of IAV. Shannon entropies, Z score, SHAPE-MaP reactivities using 1M7 and NAI, DMS-MaPseq reactivities analyzed using DREEM and shapemapper, base-pairing probabilities (*P*) for segment 2 (PB1) of A/Puerto Rico/8/1934. Reactivity values are reported as the arithmetic mean of three biological replicates. Base pairs are depicted as arcs, colored according to their probabilities. Green arcs correspond to base-pairs with *P* ≥ 0.8.

The reactivity profiles suggested that each of the eight gene segments exhibits distinct RNA conformations, as demonstrated by different patterns of reactivity profiles for each segment. Apart from the 5’ terminal end of segment 8, all gene segments at both 5’ and 3’ ends show low SHAPE reactivity, supporting the model that the 5’ and 3’ ends come together and are bound by the viral polymerase complex. We used SHAPE and DMS reactivity as constraints in RNA folding software ‘Fold’ to predict the secondary structure of each segment present in RNP complexes in virions. A maximum base pair span of 150 nucleotides (nt) was chosen to focus on local secondary structures and eliminate intra-segment long-range interactions. RNA secondary structures that were predicted in 80-100% of the SHAPE-MaP experiments with both 1M7 and NAI were considered highly confident structural elements. We identified a total of 173 predicted local RNA secondary structures; 33 in segment 1, 30 in segment 2 (**Fig 2**), 26 in segment 3, 21 in segment 4, 23 in segment 5, 17 in segment 6, 14 in segment 7, and 9 in segment 8 (**Supplementary Table 2**). Previously identified RNA structures, like the PSL2 motif and the NP-pseudoknot, were among the 173 structures that were identified (24–26).

### Selection of candidate RNA secondary structures

SHAPE-MaP and DMS-MaPseq-derived structure predictions represent the most probable structural conformation; however, many regions can adopt multiple different conformations. To identify the functionally important RNA structures, we determined base-pairing probabilities, Shannon entropy, and z-score for the entire genome. We hypothesized that functionally important structural elements, which are likely to adopt a single conformation, would exhibit high structural stability (low SHAPE and DMS reactivity) with a high base-pairing probability, low Shannon entropy, and a low z-score. Using Superfold software and SHAPE reactivity as constraints, together with a maximum base pair span parameter setting as 150, we computed base-pairing probabilities and Shannon entropy of base-pairing for each nucleotide. The Shannon entropy provides an estimate of the likelihood of a given RNA region to fold into multiple alternative conformations. ScanFold is used to calculate the z-score, which quantifies the thermodynamic stability of a region of RNA’s secondary structure, comparing it to the stability expected from a random sequence with the same nucleotide composition. We focused on the functional significance of motifs present in PB2, PB1, PA, and NP, because these gene segments and packaging signals are relatively conserved, unlike the HA and NA. We also avoided the M and NS gene segments as these transcripts are spliced, and synonymous nucleotide changes could impact RNA splicing and, therefore, the fitness of the virus. Based on these conditions, we selected a total of 15 RNA structures to assess their functional significance: 4 in PB2 (segment 1), 5 in PB1 (segment 2), 3 in PA (segment 3), and 3 in NP (segment 5) (**Fig 3A**). The selected motifs were further divided into motifs present in the canonical packaging signals (motifs A to F) (**Fig S2**) and motifs located in the central portion of the gene segments between the 3’ and 5’ packaging signal (motifs I to IX) (**Fig S3**). Motif A (PB2_2260-2275_) is a short hairpin structure present in the 3’ packaging signal of the coding region of segment 1 between positions 2260 and 2275. Motif B (PB1_40-100_) is present in the 5’ packaging signal of the coding region of segment 2 between positions 40 and 100. It is predicted to have two hairpin structures – long (stem-loop1) and short (stem-loop2) stem-loop structures with the 5’ and 3’ ends pairing to form a stem, resulting in a central 3-way junction. Motif C (PB1_2259-2284_) is a short hairpin structure present in the 3’ packaging signal of the coding region of segment 2 between positions 2259 and 2284. Motif D (PA_56-89_) is a short hairpin structure present in the 5’ packaging signal of the coding region of segment 3 between positions 56 and 89. Motif E is a 63 nt long hairpin structure (NP_1460-1522_) within the 3’ packaging signal of the coding and non-coding region of segment 5. Motif F (NP_1527-1550_) is a short hairpin structure present in the 3’ packaging signal of the noncoding region of segment 5 between positions 1527 and 1550. Motifs I to IX are predicted helical structures, between 40 and 143 nt long, and located in the central coding regions of each segment (**Fig 3A and S3**).

**Figure 3.**
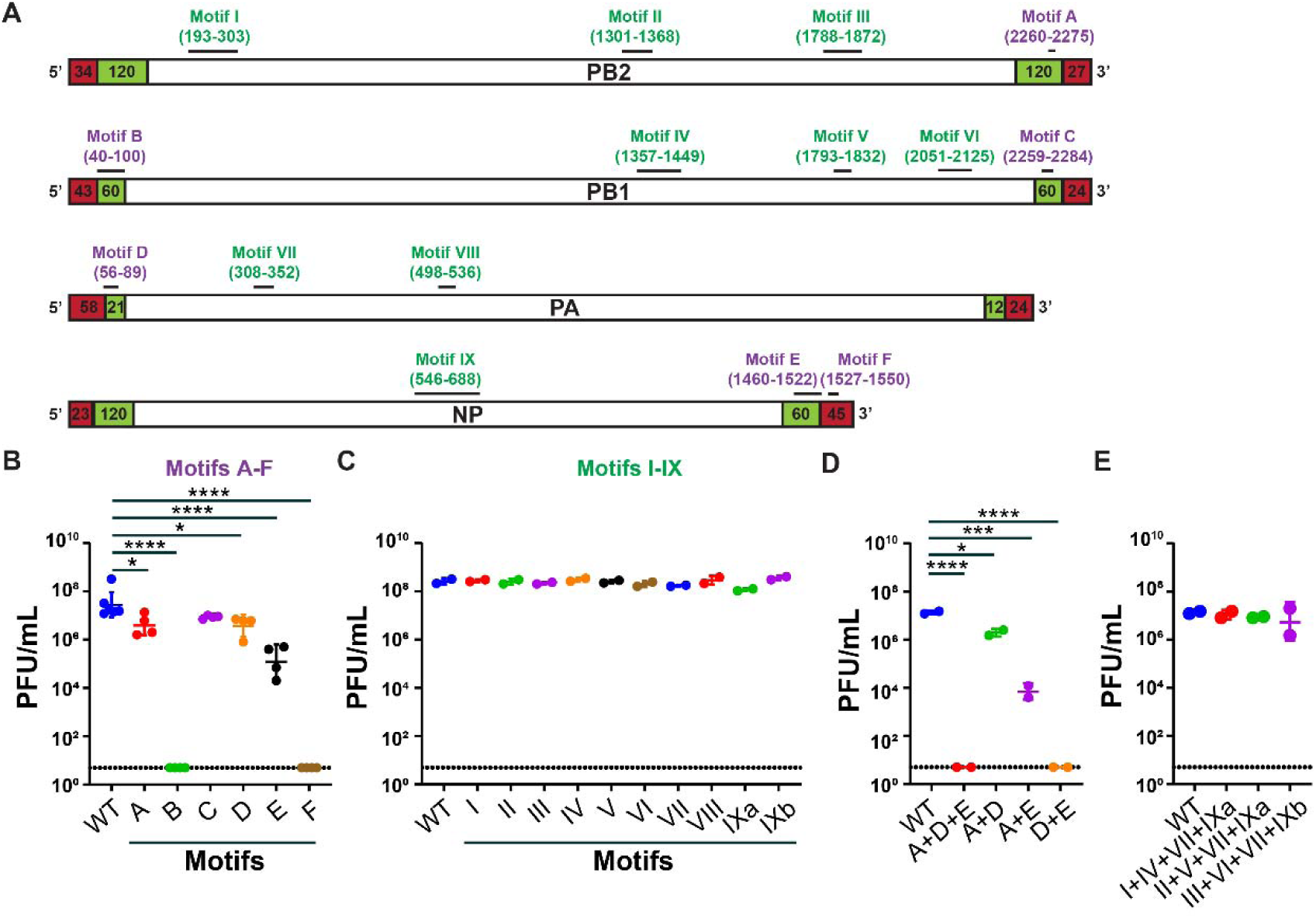
Selection and functional significance of RNA secondary structures in IAV genome. (**A**) Location of the 15 structural RNA motifs spanning both non-coding and coding regions across segments 1 (PB2), 2 (PB1), 3 (PA), and 5 (NP) of influenza A virus. Motifs located in the canonical packaging signals (highlighted by the green and red boxes at the 5’ and 3’ ends) are labelled A through F, while the ones located in the central coding region are labelled I through IX. (**B-E**) MDCK cells were infected with a low dose (multiplicity of infection of 0.001) of wild type of mutant viruses containing synonymous structural nucleotide changes in the indicated RNA motifs. Culture supernatant was collected 72 hours later, and infectious virus titer was measured by plaque assay on MDCK cells and expressed as PFU/mL. Results are the geometric mean viral titer of two to three experiments performed in duplicate. The dotted line is the limit of detection of the assay. Data was analyzed by two-way ANOVA with Dunnett’s multiple comparisons correction (\*\*\*\**p* < 0.0001, \*\*\**p* < 0.001, \*\**p* < 0.01; \**p* < 0.05).

### Functional characterization of predicted RNA secondary structures

To investigate the functional significance of selected candidate RNA structural elements, we used synonymous mutagenesis to disrupt the predicted secondary structure while preserving the encoded amino acid sequence. WT and mutant PR8 viruses were generated twice independently and expanded on MDCK cells for 72 hours (P1 stock). Following confirmation of the mutant or wild-type sequence, the infectious virus titer of the P1 stock was quantified. PR8 viruses with structural synonymous changes in motifs B (PB1_40-100_) or F (NP_1527-1550_) could not be rescued, and no infectious virus was detected upon expansion in MDCK cells (**Fig 3B**). Similarly, the virus titer of the mutant PR8 virus harboring synonymous changes in motif E (NP_1460-1522_) was reduced ∼100-fold compared to the WT virus. Finally, PR8 viruses containing structural changes in motifs A (PB2_2260-2275_) and D (PA_56-89_) had lower (∼10 fold) virus titers. Synonymous changes in motif C (PB1_2259-2284_) did not reduce the virus titer compared to WT PR8. In contrast to the synonymous RNA structural changes in the packaging signals of gene segments, synonymous changes made in motifs I to IX, did not result in lower virus titers compared to the WT PR8 virus (**Fig 3C**).

Next, we hypothesized that the effect of one RNA structural change could be enhanced by combining it with other structural changes in different segments. To test this hypothesis, we generated mutant PR8 viruses with synonymous RNA structural changes in more than one gene segment. Combining mutations in motifs A (PB2_2260-2275_), D (PA_56-89_), and E (NP_1460-1522_), which individually attenuated the PR8 virus 10-and 100-fold, did not generate a viable virus (**Fig 3D**). Similarly, combining mutations in motifs D (PA_56-89_) and E (NP_1460-1522_) also did not yield a viable virus, while a virus containing mutations in motifs A (PB2_2260-2275_) and E (NP_1460-1522_) resulted in a 1000-fold lower titer compared to the WT PR8 virus. Combining mutations in motifs A (PB2_2260-2275_) and D (PA_56-89_) did not further attenuate the virus compared to mutant A or D alone (**Fig 3D**). In contrast, combining structural mutations in motifs I (PB2_193-303_), IV (PB1_1357-1449_), VII (PA_308-352_), and IX (NP_546-688_) did not attenuate the virus compared to WT PR8 (**Fig 3E**). Similarly, combining mutations in motifs II (PB2_1301-1368_), V (PB1_1793-1832_), VIII (PA_498-536_), and IX (NP_546-688_), or III (PB2_1788-1872_), VI (PB1_2051-2125_), VII (PA_308-352_), and IX (NP_546-688_), also did not reduce the virus titer of these mutant viruses. Combined, these findings suggest that the RNA motifs present in the 5’ and 3’ packaging signals of each gene segment are significantly more important and have deleterious effects on virus replication if mutated, while synonymous structural changes in RNA motifs located in the central non-packaging region of the gene segment are not.

### Structural and functional characterization of motif B (PB1_40-100_) in the packaging signals of PR8 virus

The predicted structure at PB1_40-100_ in PR8 has not been identified or structurally characterized before. Predictions suggest that motif B (PB1_40-100_) consists of two stem-loops separated by a single residue near the 3’ end of the vRNA (**Fig 4A**). To assess whether one or both stem-loops are required for optimal replication of PR8 virus, additional mutant viruses were generated with one or more synonymous changes resulting in structural changes in stem-loop 1 or 2. Mutant virus B1 has 4 nucleotide changes that lead to the restructuring of stem-loop 1 and loss of the 3-way junction, while stem-loop 2 remains intact (**Fig 4A**). Virus B2 has an extended bulge at the 3-way junction due to impaired base pairing at the intersection, and its stem-loop 2 pairings are also compromised. Interestingly, we found that the B1 virus was attenuated ∼1000-fold, while the B2 virus grew to similar titers as WT PR8 (**Fig 4B**). To further validate this observation, we evaluated the B1 and B2 mutations in the context of other mutations in packaging signals (motif E or motifs A+D). A mutant virus containing mutations in motifs B1 and E could not be generated, while a mutant virus with mutations in motifs B2 and E had a similar virus titer compared to virus E alone (**Fig 4C**). Similarly, a virus with mutations in motifs B1, A, and D could not be generated, while a virus with mutations in B2, A, and D had an infectious titer similar to that of A+D alone. Combined, these data suggest that stem-loop 1, but not stem-loop 2, in motif B plays an important role in viral fitness. Next, single synonymous changes were introduced to differentiate which region in motif B is responsible for attenuation (**Fig 4B**). The B3 and B4 viruses contained one nucleotide mutation in the stem region of stem-loop 1, resulting in significant rearrangement of this motif, while the B5 virus was designed to expand the loop of stem-loop 1. The B6 virus harbored two nucleotide changes that led to a total loss of stem-loop 2. As a control (B7), we introduced a synonymous mutation in the loop of stem-loop 1 that did not impact the predicted RNA secondary structure (**Fig 4A**). While the virus titer of the B3 and B4 viruses was nearly 10-fold lower compared to WT PR8, the virus titer of B5, B6, and B7 was like that of WT PR8 (**Fig 4B**), indicating that the stem region of stem-loop 1 in motif B (PB1_43-93_) is critical for PR8 virus.

**Figure 4.**
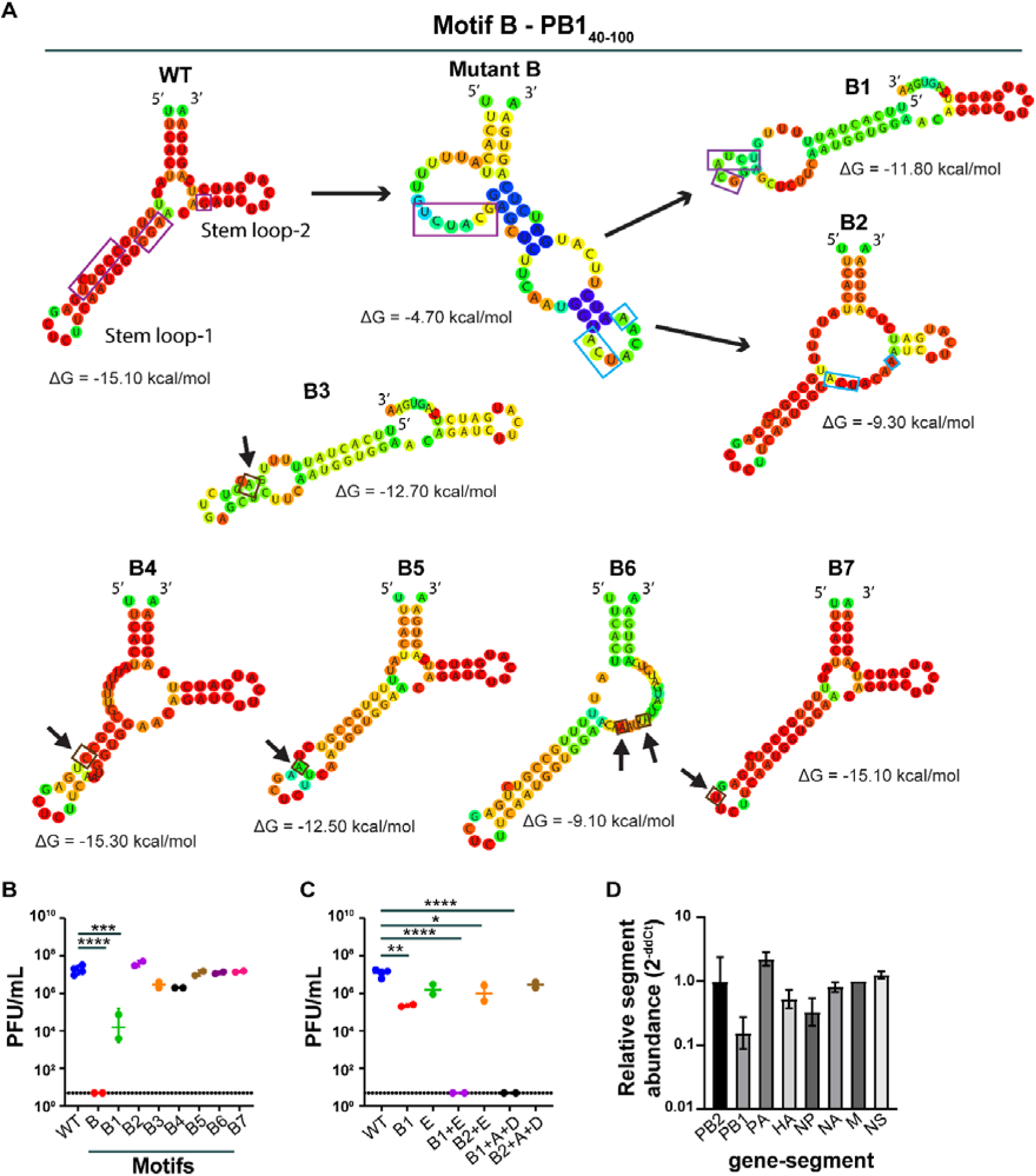
Stem region of the stem-loop 1 motif B disruption affects viral fitness. (**A**) Predicted RNA secondary structures for motif B in wild-type and mutant viruses (B1-B7). Individual residues are colored by base-pairing probabilities, according to CentroidFold. Mutations are highlighted by boxes or arrows in the predicted B1-B7 structures. For unpaired regions, the color denotes the probability of being unpaired. ΔG is the minimum free energy (MFEs) of the shown structure. (**B-C**) MDCK cells were infected with MOI = 0.001 of the indicated viruses, and culture supernatant was collected 72 hours later, then titered on MDCK cells. Bars indicate the geometric mean values of two experiments performed in duplicate, and dotted lines are the limit of detection of the assays. Data was analyzed by two-way ANOVA with Dunnett’s multiple comparisons correction (\*\*\*\**p* < 0.0001, \*\*\**p* < 0.001, \*\**p* < 0.01; \**p* < 0.05). (**D**) Relative abundance of gene segments in mutant B1 viruses. All segments were compared to segment 7 (M) vRNA and normalized to the average of WT values using the 2^-ddCt^ method. Bars represent the means of 3 independent virus preparations + standard error of the mean.

### Relative packaging efficiency of eight vRNAs

The ability of WT and mutant viruses to package all eight genome segments was assessed using a population level measure of relative vRNA segment abundance in viral particles. RNA from WT and B1 IAVs was subjected to RT-qPCR, and the abundance of segments was normalized to segment 7 (**Fig 4D**). Specific manipulation of segment 2 in the B1 virus resulted in the 6-fold reduction of packaging of segment 2 (PB1). Other than PB1, NP and HA show a 3 to 2-fold reduction, while the amount of PA segment detected was increased 2-fold.

### Motif B mutation doesn’t affect transcription and translation

To investigate whether the introduced synonymous structural mutations affect viral replication or transcription, a minigenome luciferase reporter assay was performed using a firefly luciferase reporter gene flanked by either WT or mutant (B1) 5’ and 3’ regions (residues 2318-2341 and 1-100 at the 5’ and 3’ end) of the PB1 gene segment (**Fig 5A and B**). These constructs were transfected into HEK293T cells along with plasmids expressing PB2, PB1, PA, and NP of PR8. A Renilla luciferase-expressing plasmid was included for normalization. A negative control was performed by omitting the PB1-expressing plasmid, thereby disrupting the formation of the RNA-dependent RNA polymerase (RdRP) complex (**Fig 5B**). Luciferase activity measured after 24 hours post-transfection enables us to assess viral RNA synthesis efficiency. The negative control, in which PB1 was omitted, yielded minimal signal, confirming assay specificity. No significant difference was observed between the WT and B1 mutant constructs (**Fig 5C**), indicating that the synonymous structural mutations did not affect replication or transcription of the PB1 gene segment. This suggests that motif B may instead function during later stages, such as vRNP nuclear export, intersegment interactions, genome packaging, or virion assembly, where conserved RNA secondary structures have previously been implicated.

**Figure 5.**
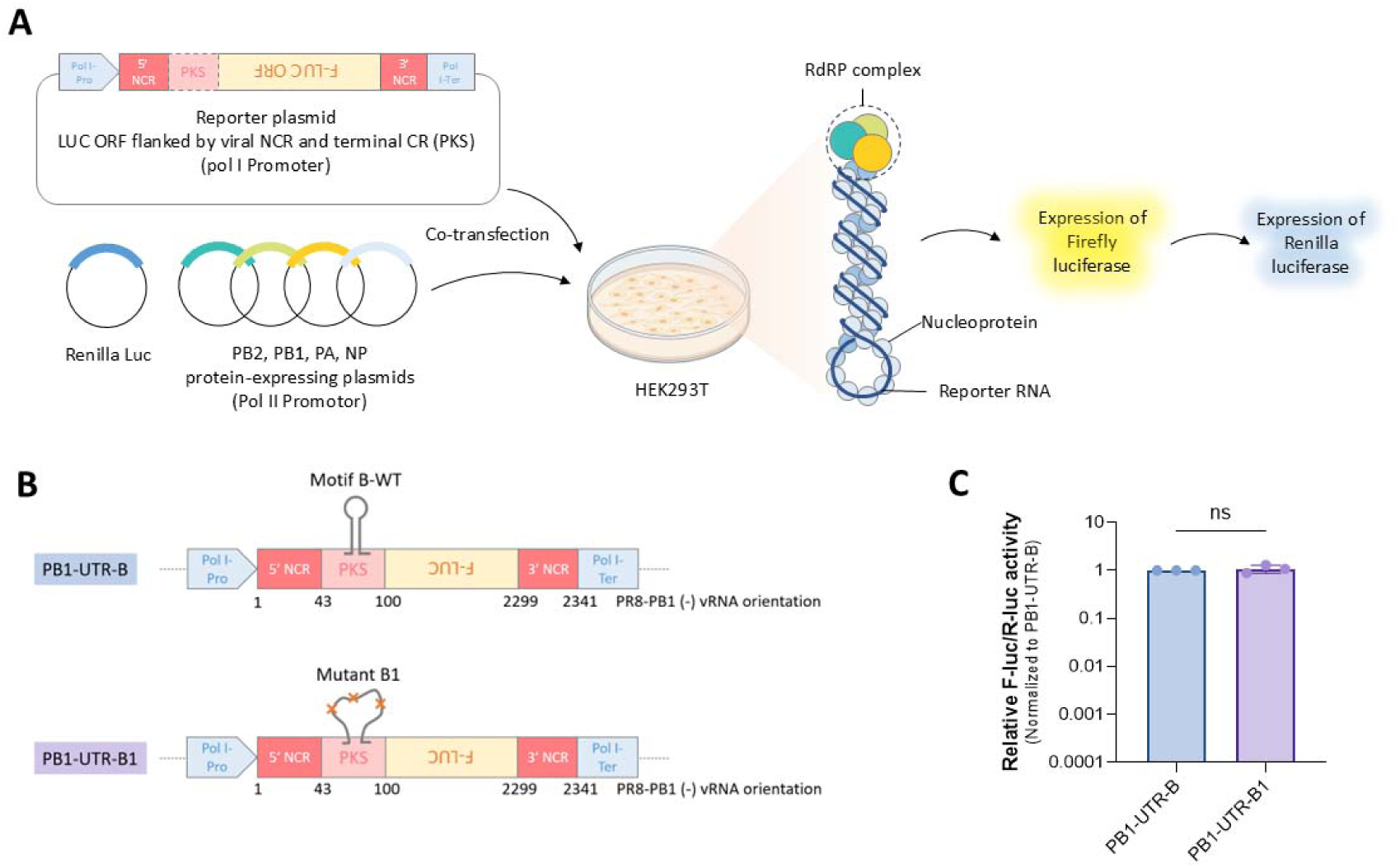
Synonymous structural changes in motif B do not affect viral polymerase activity. (**A**) Workflow of minigenome luciferase reporter assay. (**B**) Schematic representation of constructs used in the study. (**C**) Dual-luciferase reporter assay to assess viral transcription and genome replication, each combination of plasmids was assessed 3 times with corresponding to the WT-PR8 combination. Statistical analysis: Two-way ANOVA with Tukey’s multiple comparisons correction. Ns = not significant.

### Structural and functional characterization of motif E (NP_1460-1522_) and motif F (NP_1527-1550_) in the NP gene of PR8 virus

Motif E (NP_1460-1522_) is a 63-nucleotide-long hairpin structure located at the 3’ packaging signal of the NP gene segment, which has not been previously identified or structurally characterized. This region is conserved between viruses with more than 80% base pairing probability and 98.23% nucleotide conservation as reported previously by Mirska *et al*. It depicts a slightly different base pairing if the structure is predicted bioinformatically using experimentally driven data or sequence alone (**Fig 6A**). Compared to WT PR8, the viral titer of a mutant virus harboring 7 synonymous mutations in motif E was ∼100-fold reduced (**Fig 3B and 6C**), suggesting that the RNA structure is important for PR8 replication. To support this finding, a second virus was made containing a single synonymous mutation in the loop of the predicted secondary structure (E1) (**Fig 6A**). This virus was not attenuated and had similar titers compared to WT PR8 (**Fig 6C**).

**Figure 6.**
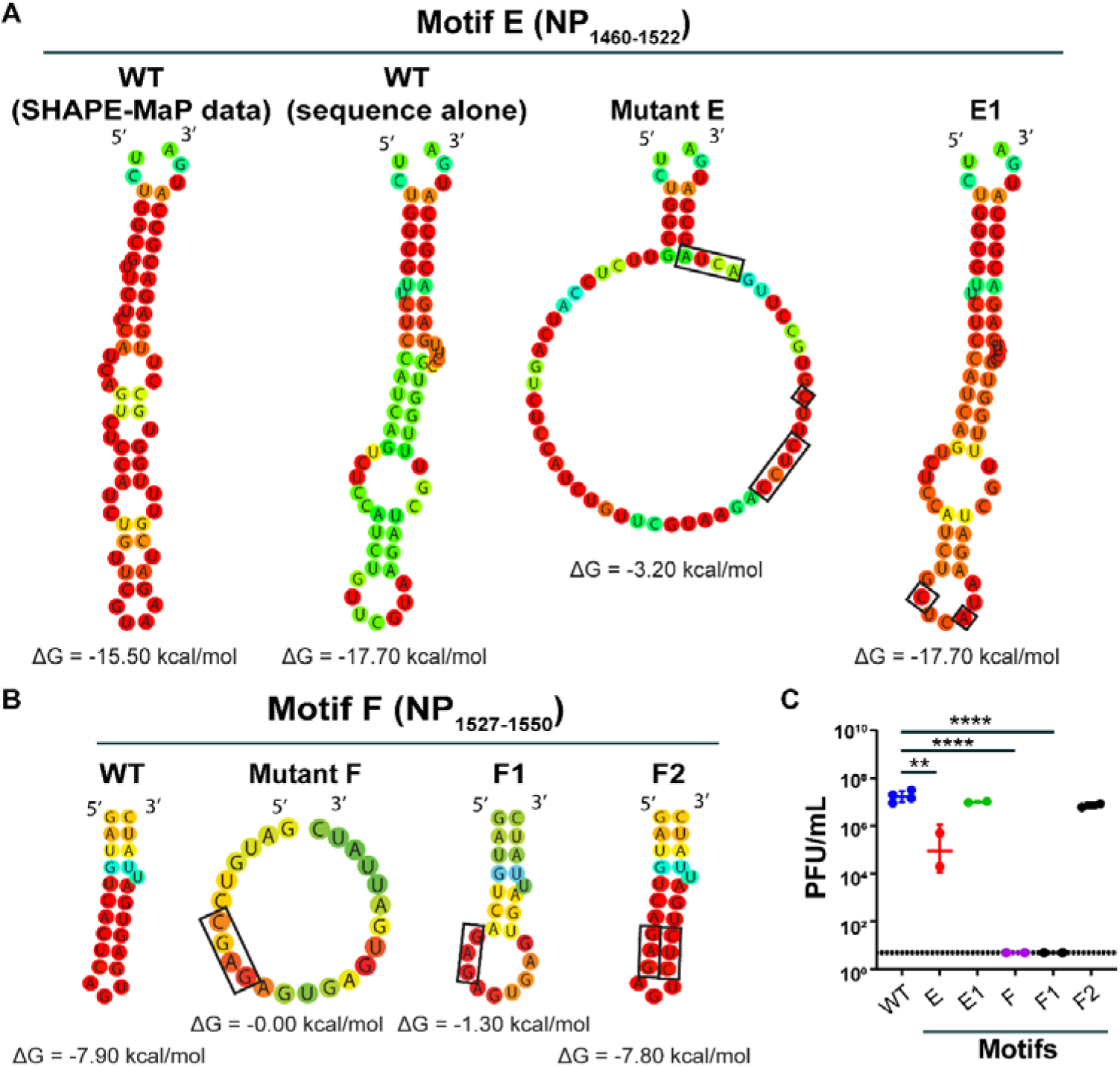
Synonymous mutational changes made in motifs E and F lead to viral attenuation. (**A-B**) Predicted RNA secondary structures for motif E and F in wild-type and mutant viruses (E, E1, F, F1, and F2). Individual residues are colored by base-pairing probabilities, according to CentroidFold. Mutations are highlighted by boxes in the predicted B1-B7 structures. For unpaired regions, the color denotes the probability of being unpaired. ΔG is the minimum free energy (MFEs) of the shown structure. (**C**) MDCK cells were infected with MOI = 0.001 of the indicated viruses, and culture supernatant was collected 72 hours later and titered on MDCK cells. Bars indicate the geometric mean values of two experiments performed in duplicate, and dotted lines are the limit of detection of the assays. Data was analyzed by two-way ANOVA with Dunnett’s multiple comparisons correction (\*\*\*\**p* < 0.0001, \*\*\**p* < 0.001, \*\**p* < 0.01).

The mutant PR8 virus containing 4 nucleotide changes in the 3’ UTR of segment 5 of the NP gene (motif F) could not be rescued following multiple independent attempts (**Fig 3B and 6C**), indicating that this short hairpin structure is essential for virus replication. To study this predicted RNA structure further, we generated two additional viruses wherein 3 nucleotides were altered, leading to a decrease in the stability of the F structure (F1) and a second virus (F2) containing compensatory mutations for the 3 nucleotide changes in F1 (**Fig 6B**). While the F1 virus could not be generated, the F2 virus grew to similar titers as WT PR8 (**Fig 6C**), suggesting that this RNA structure in the 3’ end of vRNA segment 5 is essential for virus replication or genome packaging.

## DISCUSSION

In this study, we employed high throughput *in virio* RNA probing to elucidate the RNA secondary structure landscape of the PR8 strain of IAV. This approach enabled a comprehensive mapping and identification of 173 structural elements (motifs) within all eight viral RNA (vRNA) segments under native-like conditions. Fifteen motifs in four different gene segments were selected to functionally validate their significance during virus replication. These motifs are hypothesized to act as functional RNA elements involved in genome packaging or virus replication. Synonymous mutations were introduced to destabilize the predicted RNA secondary structure, and the impact of virus replication was assessed. Synonymous mutations introduced into RNA secondary structures located within the packaging signal frequently reduced virus replication, while changes in predicted RNA secondary structures present in the central coding region did not. This finding underscores the specific importance of the packaging signals’ structural integrity for viral replication.

To establish the SHAPE-MaP method pipeline, we evaluated the amount of assay-to-assay variation, compared different propagation methods (tissue culture versus egg-grown), compared different ways of purification (ultracentrifuge and PEG), and used two different RNA-probing reagents (1M7 and NAI). Despite the NAI’s bias against guanosine and cytidine, we observed that the SHAPE reactivity was relatively consistent across different experiments and probing reagents, demonstrating a high degree of correlation (35) (**Fig S1**). We used the reactivities from each of the seven experiments to predict the secondary RNA structure of the PR8 IAV genome. The consensus secondary structure (common structure present in six out of seven experiments) was then derived from all the predicted secondary structures, which were then compared to the secondary structure predicted using DMS-MaPseq reactivity. This unprecedented amount of structural information allowed us to reliably identify 173 local RNA secondary structures in the PR8 IAV genome. The number is higher than the previously reported RNA secondary structure prediction in the IAV genome. One possible reason could be the strain of the virus. Additionally, the SHAPE protocol using both 1M7 and NAI increased our confidence in predicting more consensus structures. Another contributing factor might be our analysis pipeline that focuses on only local RNA structures. Previously, groups have reported long-range interactions or intersegment interactions (15, 16, 40). In this study, we focused on short-range or local interactions as these are easier to predict and perturb. Long-range intra-segment RNA interactions in IAV have been reported previously, but no function has been assigned to them (14). Among the 173 predicted RNA secondary structures, we identified 27 motifs located in the canonical 5’ and 3’ packaging signals, underscoring the power of structure-informed approaches in revisiting viral RNA architecture in the context of live virus and vRNPs. Out of 27, only 10 motifs were previously identified as sequence or structure by other groups in different influenza A viruses (12, 14, 15, 23, 24, 26, 30, 41–43). Functional study on 3 of these motifs, i.e., PSL2 in PB2 segment, stem-loop structure in HA segment, and a pseudoknot in NP segment, demonstrates that synonymous mutations intended to disrupt these motifs attenuated virus replication and genome packaging. To extend this observation and identify additional structural motifs packaging signals, we focused on the functional significance of 6 motifs present in the packaging signals of PB2, PB1, PA, and NP, because these gene segments and packaging signals are relatively conserved.

One of the motifs we structurally identified and characterized is motif B (PB1_40-100_), which serves as a 5’ packaging signal of the PB1 segment, comprising 2 stem-loop structures. Previous structural studies show only the stem-loop 2 conserved structure for this motif located between 79 to 93 nucleotides (14). Also, Marsh *et al* and Bolte *et al* reported that a 55 to 90 and 62 to 85 nucleotide sequence of the packaging signal in the PB1 segment is highly conserved and is important for highly efficient packaging (12, 44). We extended the findings for this motif by not only showing the structure of the whole motif but also functionally validating that a specific region, stem-loop 1, plays a crucial role in viral genome packaging. Synonymous mutations made to disrupt the stem, but not the loop, of the stem-loop 1 domain of motif B attenuated the virus ∼100-fold, while changes in stem-loop 2 did not. Currently, we do not know the function of the stem-loop 1 region and whether it is the sequence or only the structure itself that is critical for the PR8 virus. Based on our polymerase assay and segment-specific PCR, we expect that this region does not impact replication and transcription of the PB1 gene segment. Instead, the segment-specific PCR analysis suggests that this motif is critical for the packaging of the PB1 gene segment into progeny virus particles. Future studies should evaluate the role of this region in other influenza viruses, structurally characterize this stem-loop 1 region, and discern the role of structure versus sequence for the PR8 virus.

We also functionally characterized motifs E and F (NP_1460-1522_ and NP_1527-1550_) located at the 3’ end of segment 5. This motif was recently identified by Mirska *et al.* but never functionally characterized before (14, 26, 42). Synonymous mutations made to disrupt these motifs lead to a significant or total virus attenuation. The compensatory mutations made to restore the structure of NP_1527-1550_ restored the viral growth. The other motifs we chose to structurally and functionally characterize are either not studied at all or only identified as putatively important sequences (12, 14). Motif A (PB2_2260-2275_) located at the 3’ end of the PB2 segment has not been described to date. Synonymous mutations made in this motif lead to a 10-fold attenuation in a viral titer. Motif C (PB1_2259-2284_) is another motif present in the packaging signal of the PB1 segment but located at the 3’ end of vRNA. A previous study showed that nucleotide 2255 to 2304, part of motif C, is important for viral packaging (43). This motif is neither structurally identified nor functionally characterized. Motif D (PA_56-89_) is a stem-loop structure located near the 5’ end of the PA segment. It was previously identified but not functionally characterized (12, 14). Among the six motifs discussed above, synonymous structural changes in 5 of them (motif A (PB2_2260-2275_), motif B (PB1_40-100_), motif D (PA_56-89_), motif E (NP_1460-1522_), and motif F (NP_1527-1550_)) attenuated the PR8 virus. It is not clear why mutations in motif C (PB1_2259-2284_) did not attenuate the virus. Either the prediction of the RNA secondary structure is incorrect, the wrong nucleotides were mutated, or motif C does not play a significant role in virus replication and genome.

A particularly intriguing finding was the enhanced attenuation of the virus after combining some of the motifs. For example, motifs A + E or D + E resulted in a significantly greater fold reduction in virus titer compared to A, D, or E alone, suggesting a cooperative or synergistic contribution of RNA structures to viral fitness. Importantly, this additive or synergistic effect was not observed with the RNA motifs I-IX located in the central region of the gene segment. Combined, these data emphasize the importance of RNA secondary structures in genome packaging signals, but not in the more central parts of the gene segments. Our findings of enhanced attenuation of the combination also align with Bolte *et al,* which show that there is a synergy between segments formed by a redundant and plastic network of RNA-RNA interactions.

Many RNA secondary structures were identified in the non-packaging coding regions of the different gene segments of PR8. Previous studies had identified RNA-RNA interactions within these regions, suggesting an important role for these secondary RNA structures in genome packaging or the general viral life cycle (45). Extensive mutational analysis of nine different predicted RNA secondary structures, including combining mutations in different motifs (I-IX), did not significantly impair PR8 virus replication in MDCK cells. These data suggest that either the motifs are functionally redundant, their role is non-essential under the tested conditions, compensatory structural dynamics preserve their function, or our readout for attenuation is not sensitive. Alternatively, the synonymous changes that were introduced were insufficient to modify the RNA structure or impact potential RNA-RNA segment interactions. Our findings are in stark contrast with the structural mutational study done by Simon *et al.,* where they have shown that mutations disrupting a RNA structure found in the coding region of segment 4 lead to attenuation of viral titer (45).

### Limitations of our study

(1) There can be an ensemble of RNA secondary or tertiary structures, and the synonymous mutations were designed to abolish the dominant structure. It is possible that the alternative structures, for example, in the coding region or motif C, are the important structures. (2) Our bioinformatic analyses focused on local RNA secondary structures and did not allow for long-range intra-segment RNA-RNA interactions. These interactions require alternative experimental strategies, such as psoralen-based RNA crosslinking. (3) We have not tried to passage our mutant viruses to identify compensatory mutations because some mutant viruses were too attenuated or for biosafety reasons. (4) We did not perform RNA probing on vRNA and RNP complexes inside the cell. A recent study has shown that structures may fold differently in a biological context (14).

Together, these results underscore the significance of RNA secondary structure in IAV biology, offering new insights into genome organization, regulation, and the potential for structure-guided antiviral strategies. In summary, the application of SHAPE-MaP and DMS-MaPseq to H1N1 has revolutionized our understanding of non-coding structural information encoded in the viral genome. These approaches move beyond sequence analysis to a structure-function perspective, revealing that RNA-RNA interactions play a direct role in the influenza virus life cycle, from replication to immune evasion and reassortment. Future work combining RNA structure probing, functional assays, and evolutionary conservation analysis will be crucial in defining RNA-targeted antiviral strategies and in predicting the phenotypic impact of genomic changes in circulating influenza strains.

## ACKNOWLEDGEMENTS

This study was supported by grants from the National Institute of Health (R01-AI139251 to A.C.M.B) and the Children’s Discovery Institute (PD-II-2018-702 to A.C.M.B). We would like to thank Reed Trende and Kelly Pyles for helping with the analysis and other lab members of the Boon lab for their discussions. We would also like to thank Matthew Allan from Silvi Rouskin’s lab for his help setting up the DMS-MaPseq analysis pipeline.

## AUTHOR CONTRIBUTIONS

A.J. performed all the RNA probing experiments and analysis in this manuscript. A.J. and L.H. performed all the virus infection and replication studies. L.H. designed the wild-type and mutant pLuci construct and performed the polymerase assay. A.C.M.B. supervised experiments and acquired funding. A.J. and A.C.M.B. wrote the first draft of the manuscript, and all authors reviewed and edited the final version. All authors read and approved the final draft and take full responsibility for its content.

## SUPPLEMENTARY INFORMATION

**Supplementary Figure 1.**
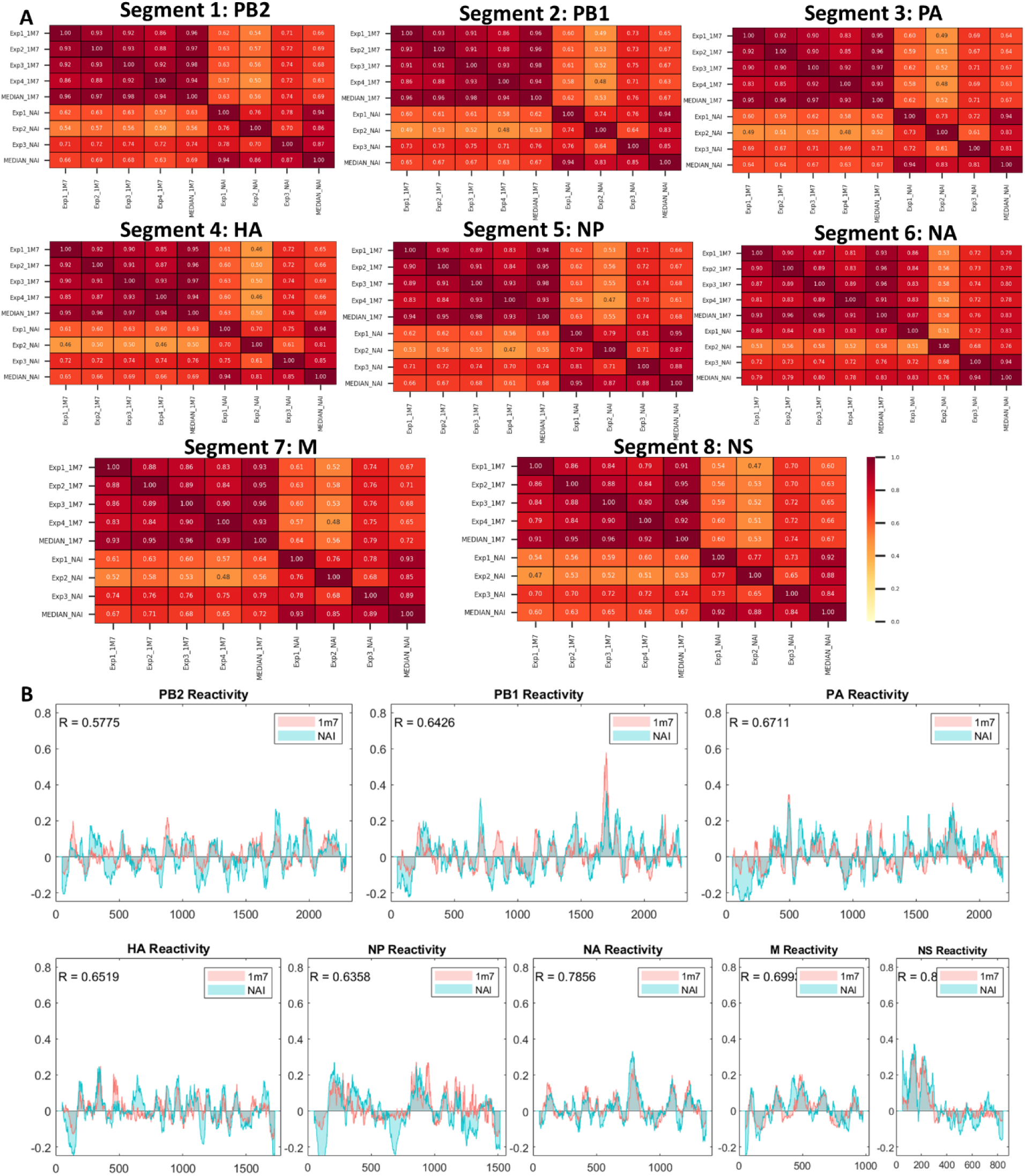
SHAPE reactivity correlation between different SHAPE reagents and replicates. (**A**) Comparison between the *in virio* SHAPE reagent (1M7 and NAI) median SHAPE reactivities of each segment. Medians were calculated over 50 nucleotide windows. Regions below zero tend to be more structured, while regions above zero indicates more flexible regions of RNA. (**B**) Pearson’s correlation of single-nucleotide SHAPE reactivities between each *in virio* replicate vRNA across independent replicates and also between the median of all replicates of 1M7 and NAI of each segment.

**Supplementary Figure 2:**
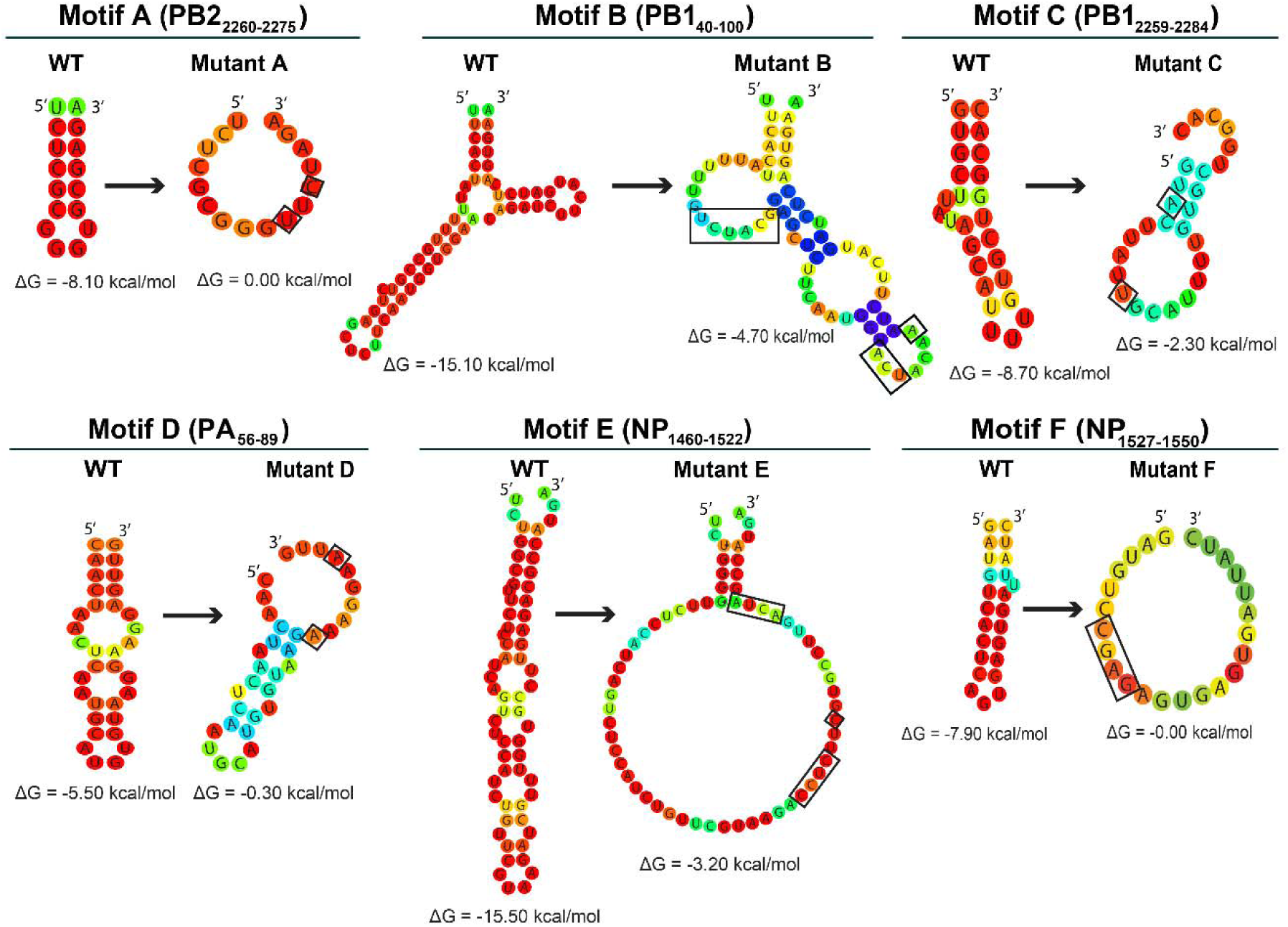
Design of the mutated structural motif in the packaging signal. Experimentally derived structure of motifs A to F, together with the predicted MFE secondary structure of the mutants. The arrow indicates where the synonymous mutations are made. Heat color gradation from blue to red represents the base pairing probability from 0 to 1. For unpaired regions, the color represents the probability of being unpaired.

**Supplementary Figure 3.**
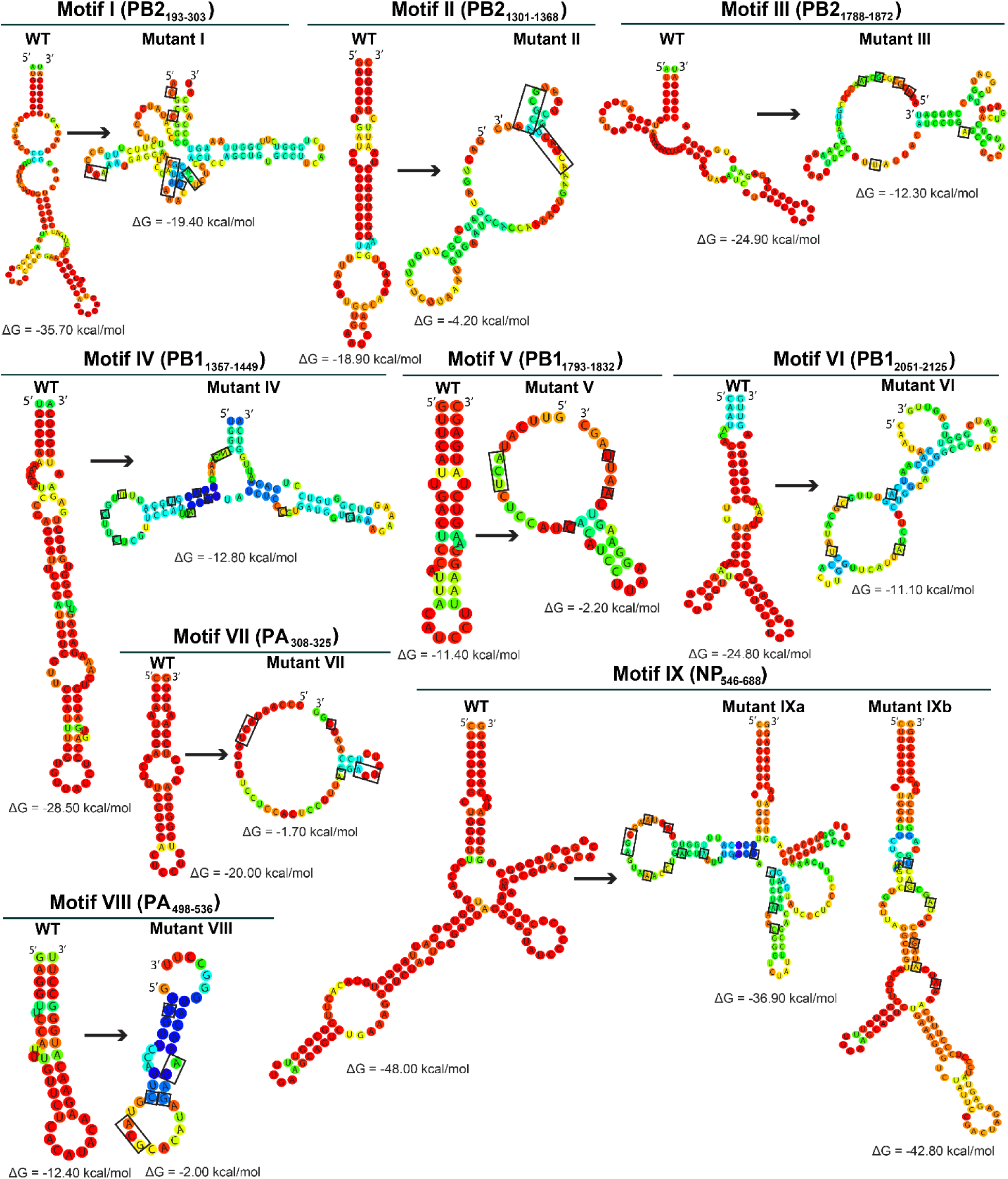
Structural predictions of wild-type and mutated motifs I-IX in PR8 virus. Experimentally derived structure of motifs I to IX, together with the predicted MFE secondary structure of the mutants. The boxes indicate where the synonymous mutations are made. Heat color gradation from blue to red represents the base pairing probability from 0 to 1. For unpaired regions, the color represents the probability of being unpaired.

**Supplementary Table 1:**
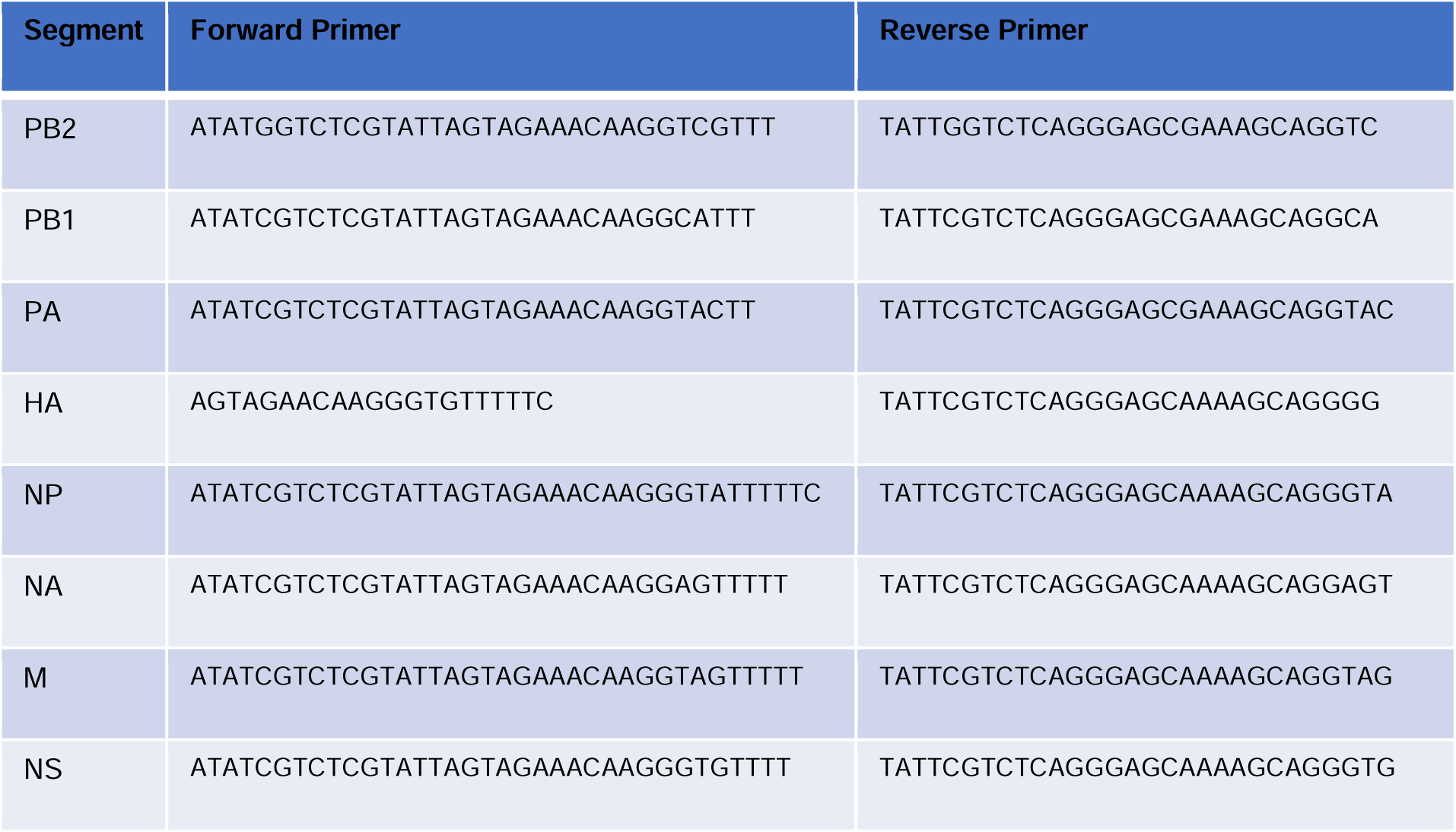
List of primer sequences used for DMS-MaPseq.

**Supplementary Table 2:**
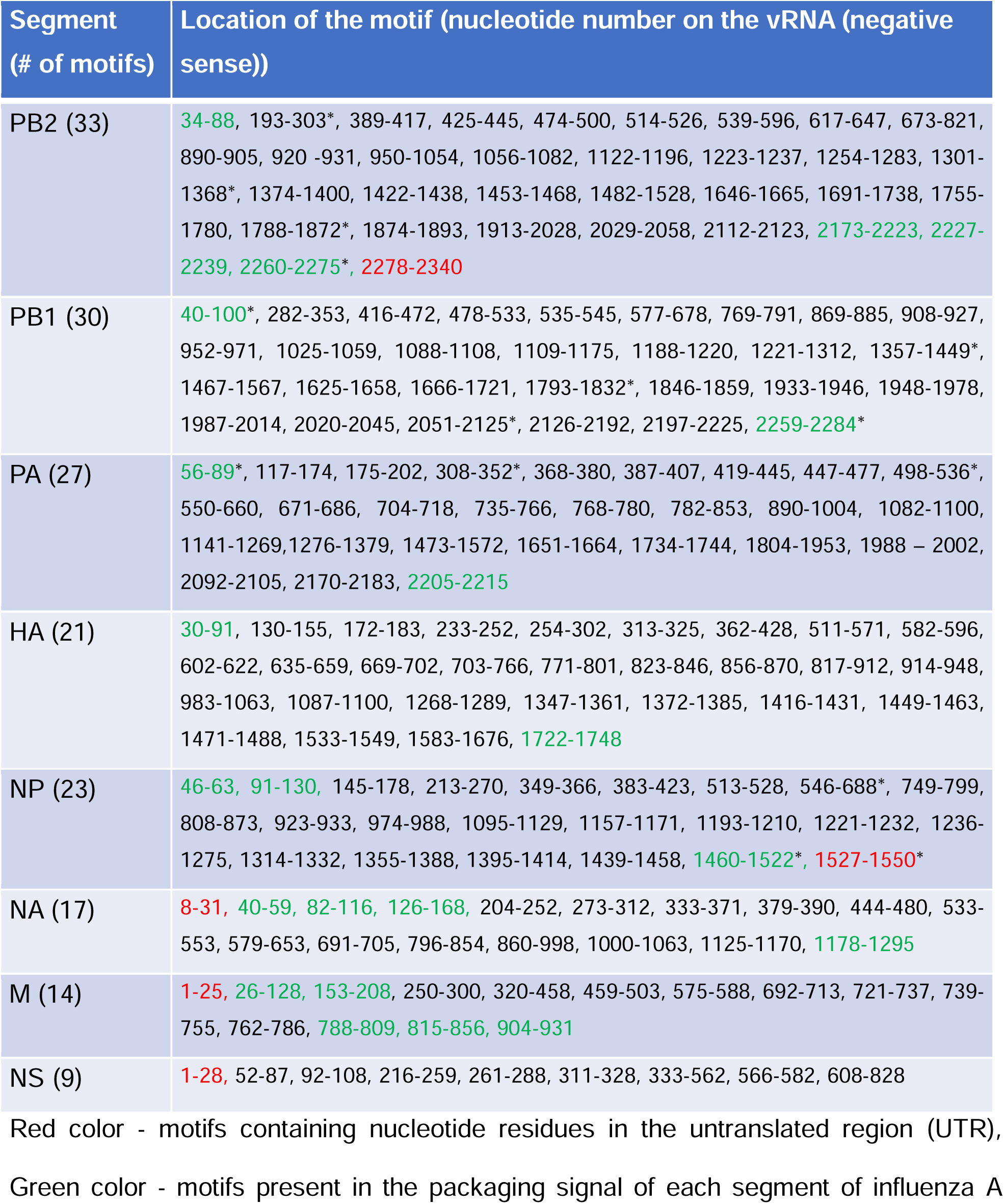
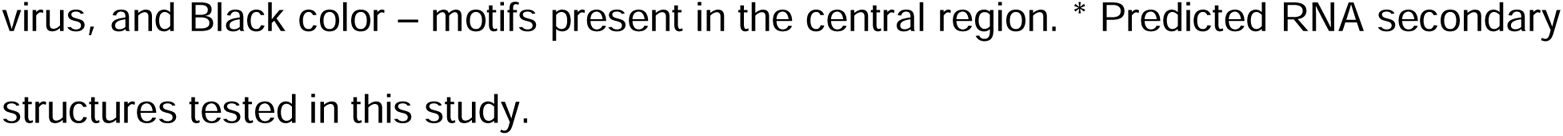
Location of all identified motifs in this study in all eight segments.

**Supplementary Table 3:**
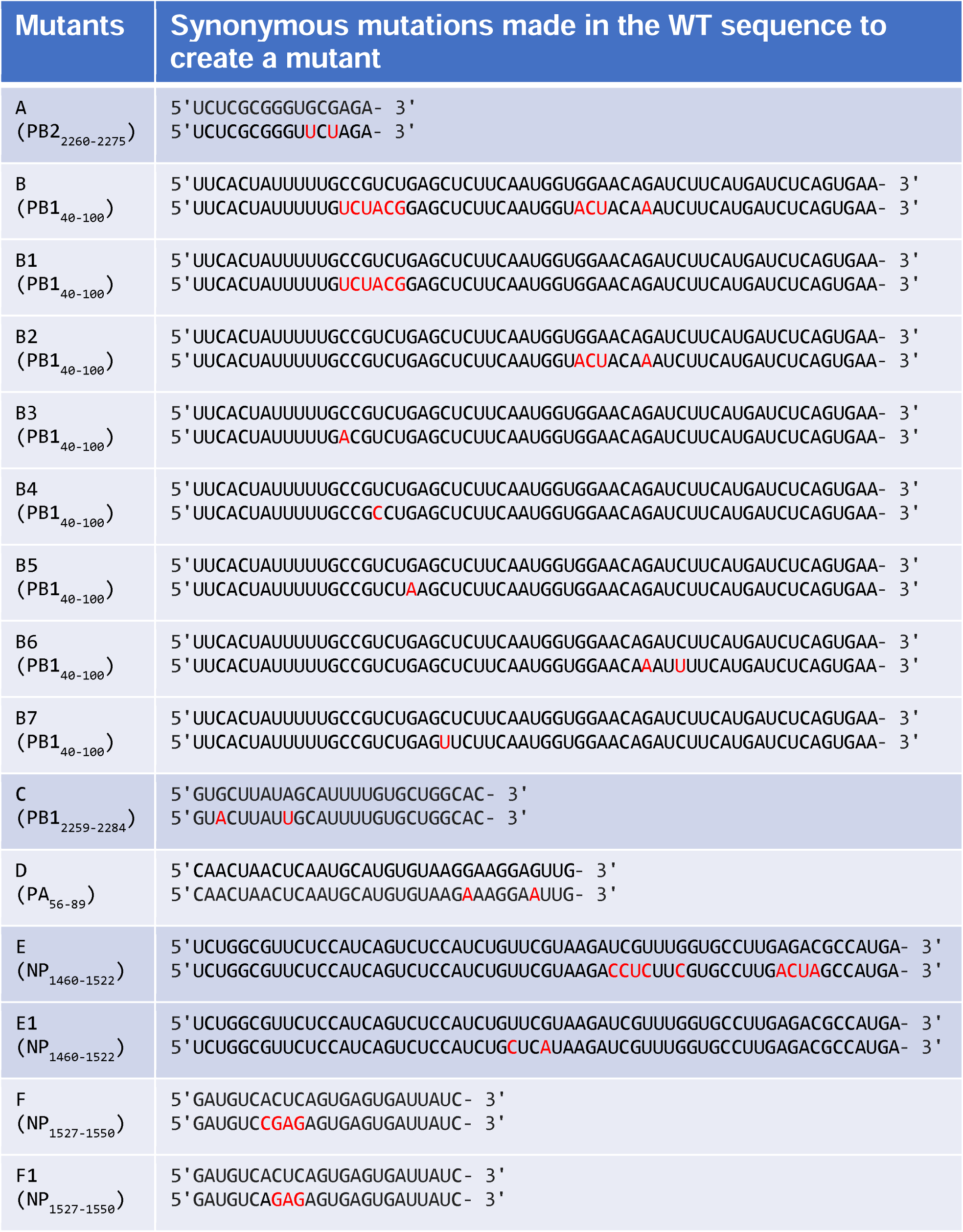
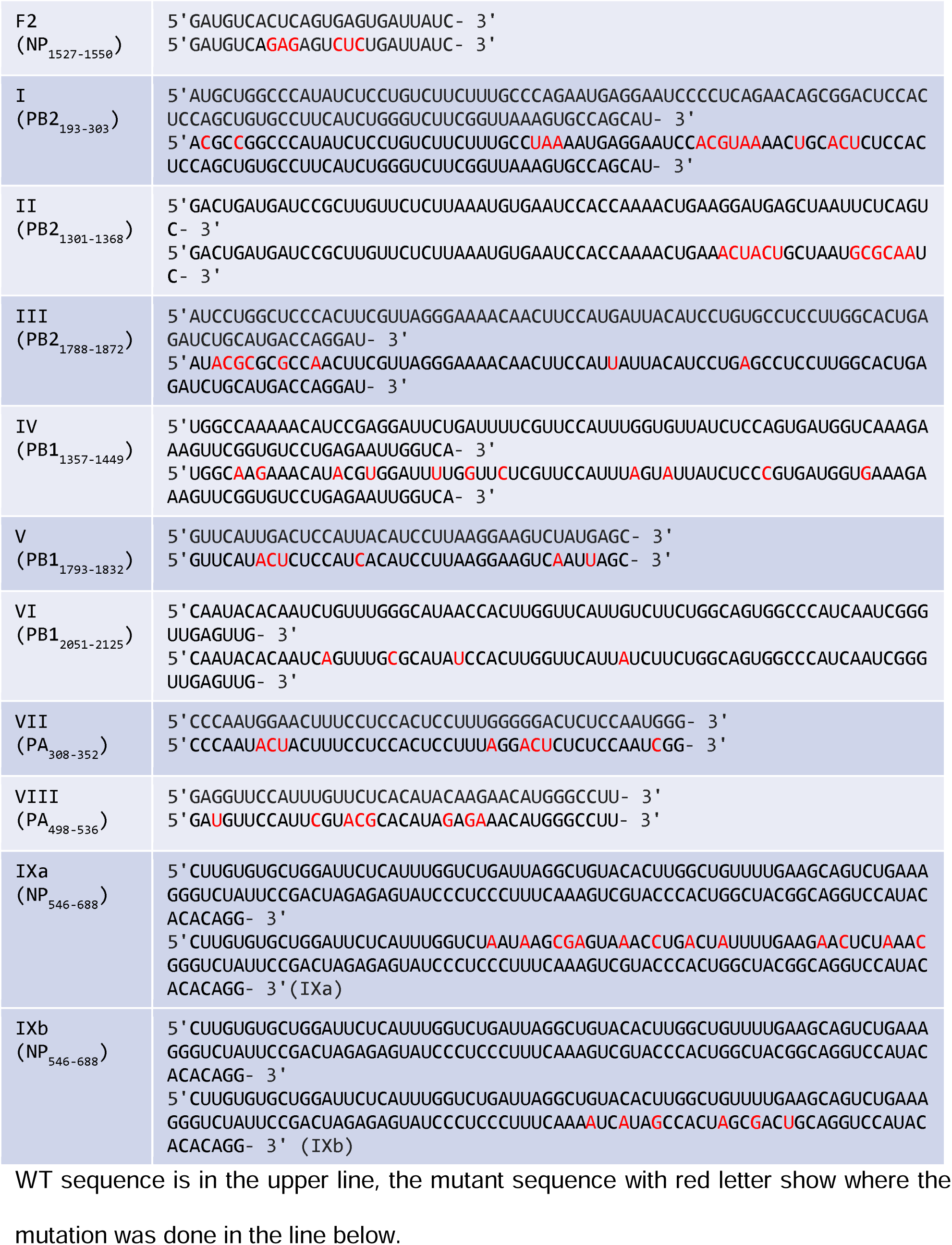
Wild-type and mutant sequence of vRNA.

